# TWISTED DWARF1 mediates myosin XI-associated vesicle trafficking required for auxin transport

**DOI:** 10.1101/2022.07.18.500530

**Authors:** Jie Liu, Jinsheng Zhu, Martin Di Donato, Pengchao Hao, Haiyun Ren, Markus Geisler

## Abstract

Defects in plant development caused by loss-of the *FKBP42, TWISTED DWARF1 (TWD1)*, have so far been accounted to a dual function of TWD1 acting as an ABCB chaperone that positively regulates ABCB biogenesis and transport activity. On the other hand, TWD1 was characterized as a modulator of actin cytoskeleton bundling and dynamics by interaction with ACTIN7, however, currently it is unclear if both events are connected.

Here, we show that TWD1 positively regulates pollen tube germination and growth by controlling actin organization. We identify and verify myosin XI-K as TWD1 interacting protein, which is most likely linking the action of TWD1 on the actin cytoskeleton. We provide evidence that myosin XI-K is required for proper auxin exporter trafficking and auxin export. Further, we show that ER-localized TWD1 reshapes the ER network to overlay actin cables similar to mutations of myosin-XI and thus controls cytoplasmic streaming.

In summary, our data support a model in that TWD1 functions as an ER–actin adapter proteins involved in myosin-dependent ER motility and cargo trafficking. Our findings provide a molecular explanation for the defects in early ABCB biogenesis in *twd1* that are caused by defects in the three-way interaction between the ER, cytosolic myosin-XI and F-actin.

## INTRODUCTION

A remarkable feature of plant cells is the extensive motility of organelles and the cytosol, which was originally defined as cytoplasmic streaming first described in 1774 (Corti, 1774). Pharmacological experiments suggested early on that the cytoskeleton plays an important role during cytoplasmic streaming, which is caused by tubule extension of the endoplasmic reticulum (ER) network, found to be much more dynamic in plants than in animals. Also unlike in animal cells, in plant and yeast cells the cortical ER network, which is defined as ER directly beneath the PM, overlies the actin cytoskeleton instead the microtubules (MT) and is mainly used to remodel the ER (Sparkes et al., 2009; Sparkes et al., 2009).

Cytoplasmic bulk flow in the cell is thought to be caused by unidirectional actin filament bundles and organelle-associated myosin XIs, a plant-specific class of myosin motor proteins (Shimmen and Yokota, 2004; Tominaga et al., 2013). Previously, it was demonstrated that ER dynamics are driven primarily by the ER-associated myosin XI-K and that myosin XI deficiency affects organization of the ER network and orientation of the actin filament bundles (Ueda et al., 2010). Thus, dynamic three-way interactions between ER, F-actin, and myosin XIs seem to determine the architecture and movement patterns of thick ER strands, and cause cytosol hauling traditionally defined as cytoplasmic streaming (Ueda et al., 2010; Ueda et al., 2015).

Plant myosins that are split into two classes, myosins VIII and XI, have recently received much attention. In Arabidopsis, the myosin XI family has thirteen members and is expanded in comparison to lower plants and most members display some form of motor activity implicated in cytoplasmic streaming, actin organization and plant development (Duan and Tominaga, 2018). Investigation of myosin loss-of-function plants demonstrated that myosin XI-K is a principal driver of cytoplasmic streaming and organelle trafficking, with myosins XI-1 and XI-2 also contributing to these processes (Prokhnevsky et al., 2008; Peremyslov et al., 2010; Abu-Abied et al., 2018). Recently, double mutant analyses revealed overlapping functions of the four class XI myosins, with *XI-K XI-1* loss-of-function plants showing the largest reductions in organelle velocities corelating with stunted plant growth (Prokhnevsky et al., 2008).

Although the three-way interaction between the ER, cytosolic myosin-XI, and F-actin is widely accepted as cause of ER streaming (Ueda et al., 2010), the mechanisms underlying stable interaction of the ER membrane with actin are unknown. While early electron microscopy studies suggested a direct attachment of the plant ER with actin filaments, it is today thought that yet-unknown proteins facilitate anchoring of the ER membrane with the actin cytoskeleton. Recently, the plant-unique SNARE protein, SYP73, which contains a transmembrane domain targeted to the ER, was demonstrated to acts as ER membrane-associated actin-binding protein (Cao et al., 2016). Over-expression of SYP73 causes a rearrangement of the ER over actin and, similar to mutations of myosin-XI (Prokhnevsky et al., 2008; Peremyslov et al., 2010; Abu-Abied et al., 2018), loss of *SYP73* reduces ER streaming and affects overall ER network morphology and plant growth (Cao et al., 2016). Moreover, NET3B, a member of the NETWORKED (NET) super family, was also suggested to regulate ER-actin interaction: overexpression of *NET3B* enhances the interaction between the ER network and the actin cytoskeleton leading to a higher percentage of co-alignment between these two structures thought to slow-down ER membrane diffusion (Wang and Hussey, 2017). The current picture that emerges is that these ER–actin adapter proteins modulate ER structure and movement by working in conjunction with myosin XI proteins. The adapter proteins would provide an anchorage that brings ER and F-actin closer together allowing the myosin driven movement to take place.

The functional three-way interaction between the ER, myosin-XI and F-actin as a driving force for plant development seems to be indirectly and directly connected with the action of the plant hormone, auxin. Application of IAA (3-indolyl acetic acid), the most prominent auxin, accelerates cytoplasmic streaming at low concentration and inhibits it at high concentration in several plant cell types (Sweeney and Thimann, 1942). The effect of auxin on cytoplasmic streaming is at least partially caused by the effect of auxin on the actin cytoskeleton. On one hand, auxin induces actin gene expression: *ACTIN7*, one of the three vegetative actin isoforms in Arabidopsis, responds strongly to auxin and other environmental stimuli (Zhu and Geisler, 2015). On the other, auxins (and auxin transport inhibitors) were shown to affect actin bundling, however, most likely in an indirect fashion (Dhonukshe et al., 2008). Thus, auxin action on actin requires the action of an integrating protein, the identity of which is unknown (Zhu and Geisler, 2015). Such auxin-actin integrators were suggested to be in fact part of the auxin export complex itself, which is based on the fact that some of these proteins bind the non-competitive auxin efflux inhibitor, NPA (1-*N*-naphtylphtalamic acid), itself known to alter actin bundling (Zhu and Geisler, 2015). A valid integrator candidate is the ER and PM membrane-associated FKBP, TWISTED DWARF1 (TWD1), which owns a dual function as a chaperon of ABCB-type auxin exporters (Zhu and Geisler, 2015; Zhu et al., 2016). First, TWD1 controls early ABCB biogenesis underlined by the finding that a subset of auxin-transporting ABCBs (ATAs) are retained on the ER in *twd1* (Wu et al., 2010; Wang et al., 2013). Second, TWD1 regulates ABCB auxin transport activity on the PM most likely by a hidden PPIase refolding a D/E-P motif on ABCBs that is required for auxin transport (Geisler and Hegedus, 2020; Hao et al., 2020). On the other hand, TWD1 was shown to physically interact with ACTIN7 and determine actin filament organization and dynamics (Zhu et al., 2016). Moreover, TWD1 is required for NPA and NSAID-mediated actin cytoskeleton remodeling (Zhu et al., 2016; Tan et al., 2020). As such, the TWD1-ACTIN7 axis seems to control the plasma membrane presence of transport proteins, however, while loss of *ACTIN7* function seems to effect PM proteins in general (Mao et al., 2016; Zhu et al., 2016), TWD1 seems to be specific for a few ATAs (Wu et al., 2010; Wang et al., 2013).

Currently it is unclear if the function of TWD1 as an ABCB chaperon and on actin cytoskeleton remodeling are connected and if the latter is direct and/or caused via auxin. Support for a direct regulatory involvement of the auxin efflux machinery in cytoplasmic streaming of plant cells was provided by the finding that cytoplasmic steaming was drastically impaired by depletion of *ABCB19* (Okamoto et al., 2016). This is surprising, as until now, no components other than the actin-myosin XIs and ER-related proteins were reported as being involved in cytoplasmic streaming.

Here, we demonstrate that TWD1 plays a key role in actin organization and tip-directed vesicle transport in pollen tubes in an action that is widely independent of ABCBs. We show that over-expression of *TWD1* causes a striking rearrangement of the ER over actin similar to mutations of myosin-XI (Peremyslov et al., 2010; Ueda et al., 2010), while loss-of-*TWD1* reduces significantly myosin-dependent ER motility and cargo trafficking. Our data imply a model in that the dynamic rearrangement and streaming of the ER network depends on an interaction between TWD1 and myosin XI-K bridging the ER membrane with actin filaments.

## RESULTS

### TWD1 plays a key role in pollen germination and pollen tube tip growth

Previous studies have shown that TWD1 is strongly expressed in the tapetum and meiocytes of Arabidopsis stamen and that TWD1 is involved in pollen maturation by differentially activating ABCB1 and ABCB19 leading to altered auxin transport during stamen development (Liu et al., 2022). The number and viability of *twd1* pollen was significantly reduced (Liu et al., 2022), however, the impact of TWD1 on pollen physiology had not been tested yet.

Pollen was chosen because as a single cell system, pollen auxin homeostasis is not dependent on external sources. Furthermore, in contrast to other pollen-specific auxin transporters, such as PIN8 and PILS5 (Feraru et al., 2012), ABCB1, ABCB19 and ABCB4 are only marginally expressed in pollen (Supplementary Figure 1A) allowing for a dissection of ABCB-dependent and independent actions of TWD1. To deeper investigate the effect of TWD1 on pollen physiology, we further assessed the expression of *TWD1* in mature pollen. qRT-PCR analysis indicated that *TWD1* was moderately expressed in pollen grains but that expression of *TWD1* increased significantly during pollen germination after 6h of incubation in germination medium compared to a hydration of 0.5h (Figure 1A).

**Fig 1:**
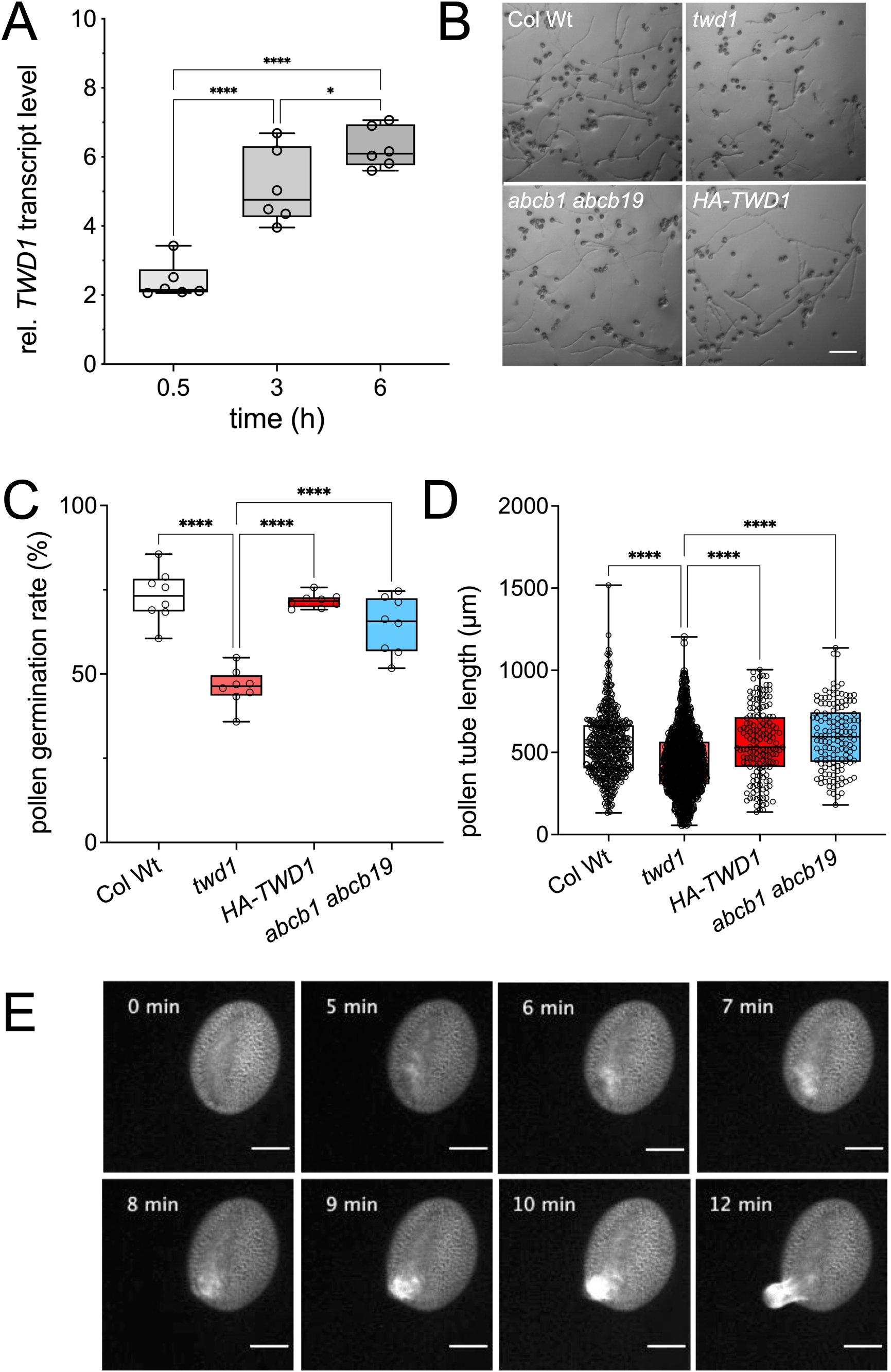
TWD1 is expressed in germinating pollen and pollen tubes and regulates pollen tube germination and growth. **A:** Quantitative real-time PCR analysis of *TWD1* expression in germinating pollen analyzed at 0.5 h, 3 h and 6 h after hydration. Significant differences (Ordinary one-way ANOVA) ± SE (n =6) of *TWD1* expression are indicated: *, *p* < 0.05; **, *p* < 0.01; ***, *p* < 0.001; ****, *p* < 0.0001. **B:** *In vitro*-germinated pollen tubes from wild-type (Wt; Col-0), *twd1* (*twd1-3*), *abcb1 abcb19 (abcb1-100 abcb19-101)*, and *HA-TWD1* (*35S:HA-TWD1* in *twd1-3*) pollen 6h after germination. Bar = 150μm. **C:** Germination frequency of Wt (Col-0), *twd1* (*twd1-3), abcb1 abcb19 (abcb1-100 abcb19-101)*, and *HA-TWD1* (*35S:HA-TWD1* in *twd1-3*) pollen 6h after initiation of germination. Significant differences (Ordinary one-way ANOVA) ± SE (n = 7) in germination frequencies are indicated: *, *p* < 0.05; **, *p* < 0.01; ***, *p* < 0.001; ****, *p* < 0.0001. **D**: Pollen tube lengths of germinated wild-type (Col-0), *twd1* (*twd1-3*), *abcb1 abcb19 (abcb1-100 abcb19-101)* and *HA-TWD* (*35S:HA-TWD1* in *twd1-3*) pollen were analyzed 6h after germination. Between 150 and 1000 pollen from three independent experiments were analysed. Significant differences (Ordinary one-way ANOVA) ± SE (n = 7) in tube lengths are indicated: *, *p* < 0.05; **, *p* < 0.01; ***, *p* < 0.001; ****, *p* < 0.0001. **E**: TWD1-CFP (*TWD1:TWD1-CFP*) is expressed in the cytoplasm of ungerminated pollen grains and at the germination site just prior to pollen tube germination. After germination, the TWD1-CFP signal is restricted to the subapical domain during pollen tube growth; bar = 10 μm. Supplementary information can be found in Suppl. Fig. S1 and Suppl. Movie S1.

To investigate the function of TWD1 during pollen germination and pollen tube growth, we quantified *in vitro* pollen germination of wild-type (Wt; Col-0), *twd1* (*twd1-3*) and *abcb1 abcb19 (abcb1-100 and abcb19-101)*; the two latter showing a remarkable similar mutant phenotype (Geisler et al., 2003). At 6h post germination, the mean pollen germination frequency of wild-type was 73.19% ± 2.7, while that of *twd1-3* was 46.1% ± 1.9, indicating that loss of *TWD1* resulted in a significant decrease (roughly 40%) in germination frequency (Figures 1B and 1C). This defective pollen germination phenotype could be fully restored by expressing *HA-TWD1* under the strong, constitutive *CaMV35S* (*35S*) promoter (71.7% ± 0.7) in the *twd1-3* background.

Next, we compared the mean lengths of Wt and *twd1-3* pollen tubes at 6h after *in vitro* germination *twd1-3* pollen tubes (444.5 ± 190 um at 6h) were significantly shorter than those in Wt plants (543.0 ± 190 um at 6h; Figures 1B and 1D).

Interestingly, there was no significant difference in germination frequency and pollen tube growth between Wt and *abcb1 abcb19*. These results demonstrate that loss-of-TWD1 function leads to a reduction in pollen tube germination and growth in an action that is independent of ABCB1 and ABCB19-mediated auxin transport.

To further investigate the function of TWD1 in pollen germination and pollen tube tip growth, time-lapse imaging of TWD1-CFP expressed under the control of its native promoter (*TWD1:TWD1-CFP*) in *twd1* (*twd1-1*) pollen (Wu et al., 2010) was performed (Figure 1E; Supplemental Figure 1B-C and Movie 1). Prior to germination, TWD1-CFP was observed in the pollen cytoplasm (Supplemental Figure 1B). With the begin of pollen germination, TWD1-CFP was observed as a ‘collar-like’ structure at the site of germination (Figure 1E; Supplemental Figure 1C-D). Previous reports have shown that the actin cytoskeleton shows a similar structure at the site of pollen germination (Pierson and Cresti, 1992; Gibbon et al., 1999; Wu et al., 2010; Liu et al., 2018). After germination, the TWD1-CFP signal was concentrated in the subapical region of the pollen tube (Fig. 1E; Supplemental Figure 1C and Movie 1) that contains a highly dynamic “actin fringe” structure as well as a peak region for exocytosis. All the above results suggest a functional correlation between TWD1 and the actin cytoskeleton during pollen germination.

### TWD1-loss-of-function alters actin organization and tip-directed vesicle transport in pollen tubes

To further investigate a functional correlation between TWD1 and the actin cytoskeleton, we generated transgenic plants co-expressing *TWD1-mCherry* and *lifeact-mEGFP* (Riedl et al., 2008), both under the control of the pollen-specific promoter, *Lat52* (Twell et al., 1989). In the germinating pollen grains, we observed colocalization of TWD1-mCherry with lifeact-mEGFP near the germination site (Figure 2A), suggesting that TWD1 and actin are jointly involved in high-density vesicle transport at the pollen germination site.

**Fig. 2:**
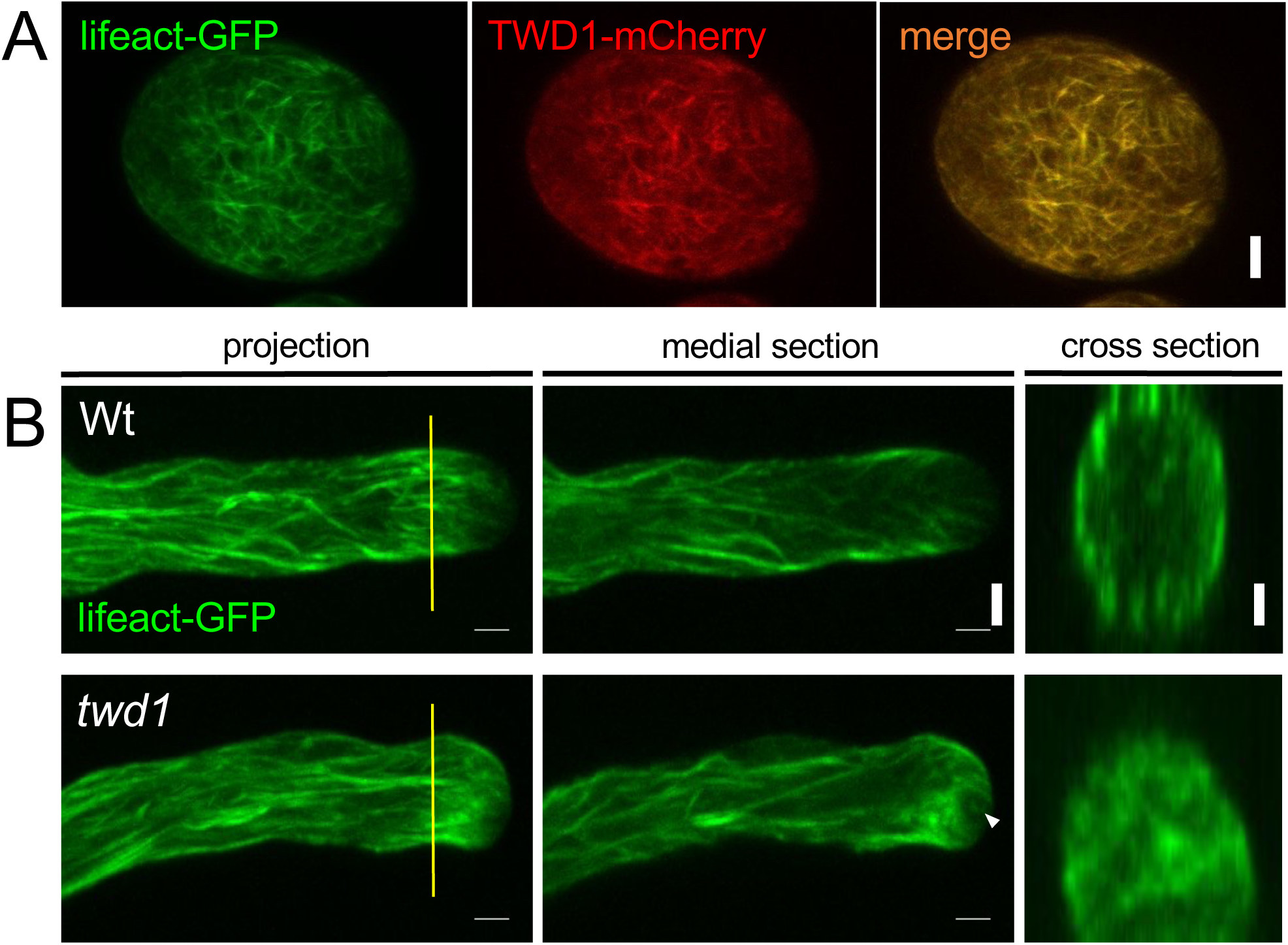
TWD1 controls pollen germination and pollen tube growth by affecting actin organization. **A:** TWD1 (*Lat52:TWD1-mCherry*) co-localizes with actin bundles (*Lat52:lifeact-GFP*) mainly in the region of the germination site; bar, 10 μm. **B:** Subapical actin filaments are disorganized in *twd1* (*twd1-3*) pollen tubes. Sites of optical sections leading to cross sections are indicated by yellow lines; bar, 10 μm.

Next, we examined the effect of TWD1 on the actin cytoskeleton at the pollen tube tip. The subapical region of Wt pollen tubes reveals a so-called actin fringe structure that is characterized by short parallel actin cables (Figure 2B), while most of the underlying actin filaments aligned longitudinally (Lovy-Wheeler et al., 2005). However, in the *twd1-3* pollen tube the F-actin became obviously disorganized within the apical-subapical region (Figures 2B). Cross sectioning revealed that the subapical actin filaments are no longer concentrated in the cortex of the pollen tube, suggesting that TWD1 is required for actin fringe formation.

### TWD1 remodels the ER network to overlay actin cables but does not bind actin directly

To be better able to quantitatively investigate the effect of TWD1 on the organization of the actin cytoskeleton, we analyzed the behavior of the actin marker, fABD2-GFP (Ye et al., 2009; Montes-Rodriguez and Kost, 2017) in the presence and absence of TWD1-mCherry in tobacco epidermal cells after agrobacterium-mediated co-transfection. Interestingly, we found that in cells expressing low levels of TWD1-mCherry, the ER network partially overlapped with the actin cytoskeleton (Figures 3A and 3B). However, in cells expressing high levels of TWD1-mCherry, the ER network almost completely overlapped with the actin cytoskeleton cables (Figures 3A and 3B), indicating that TWD1 as major ER resident (Suppl. Figure S2A) has the capacity to remodel the ER network leading to an overlay with actin cables. This ER remodeling capacity of TWD1 is most likely independent of the microtubule network because co-expression of TWD1-mCherry with microtubule marker, TUA6-GFP, indicated that the modified ER network labeled by TWD1-mCherry does not overlap with the microtubule cytoskeleton (Suppl. Figure 2B). Moreover, high expression of PIN6-GFP together with the ER lumen marker, ERYK, did not lead to ER rearrangement resembling an actin-like organization, which supports the specificity of the ER shaping effect of TWD1 (Suppl. Figure S2C).

**Fig. 3:**
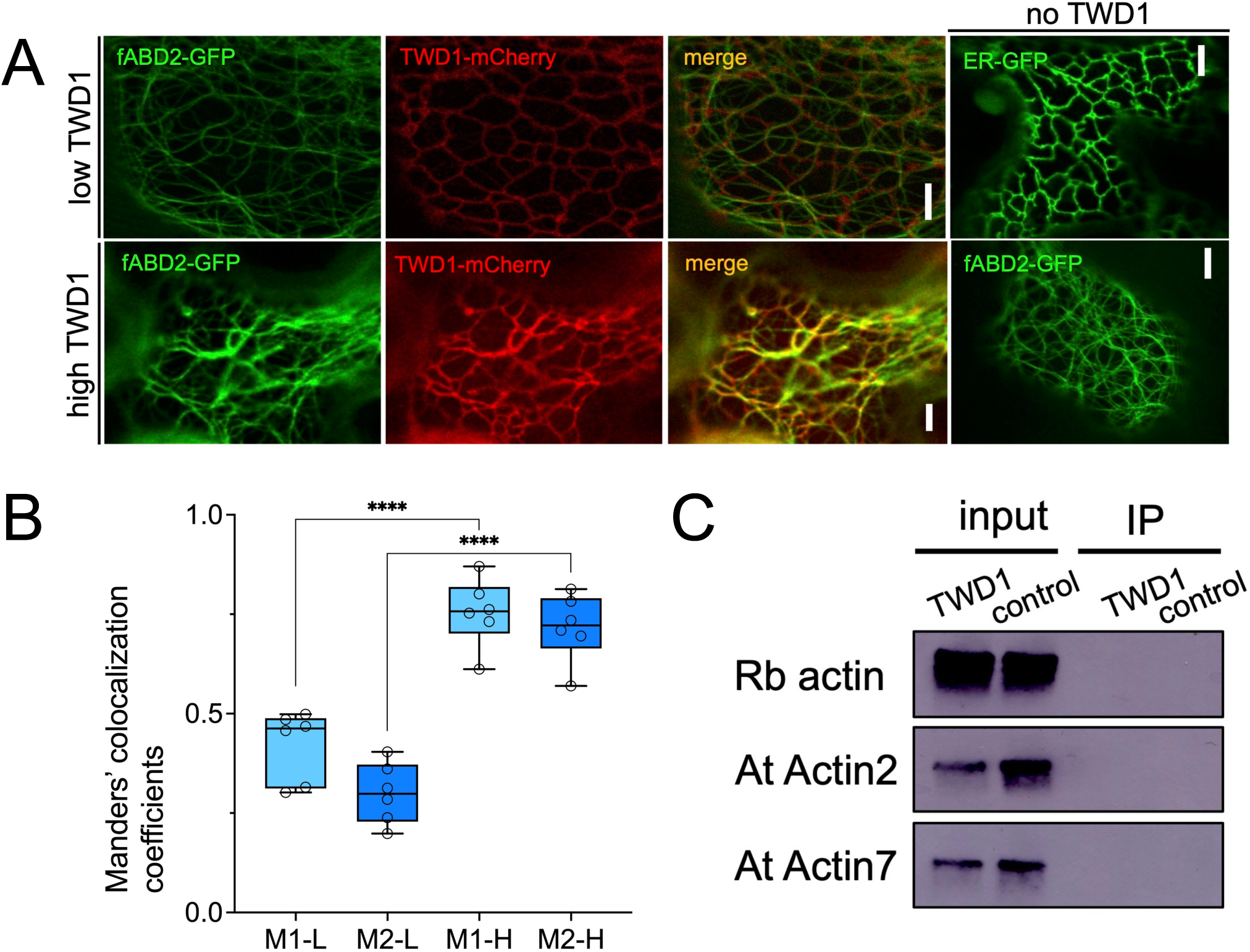
TWD1 remodels the ER over the actin cytoskeleton but does not bind actin directly. **A:** Confocal images of tobacco leaf epidermal cells expressing the actin marker, fABD2-GFP (*35S:fABD2-GFP*), and TWD1-mCherry (*35S:TWD1-mCherry*). In cells expressing TWD1-mCherry at low levels (low TWD1), the ER network is only partially overlapping with the actin cytoskeleton, while in cells expressing higher levels of TWD1-mCherry (high TWD1), the ER network is modified and overlaps almost completely with the actin cytoskeleton cables. ER-GFP (*35S:GFP-HDEL*) and fABD2-GFP (*35S:fABD2-GFP*) in the absence of TWD1 (no TWD1) are given as reference. **B:** Colocalization (Manders’ colocalization coefficients (MCC) between fABD2-GFP and TWD1-mCherry indicate that in cells expressing high levels of TWD1-mCherry (high), fABD2-GFP and TWD1-mCherry largely overlap and that the degree of overlap was significantly higher than in cells expressing low levels of TWD1-mCherry (low). M1 indicates the fraction of fABD2-GFP fluorescence overlapping with TWD1-mCherry fluorescence and M2 indicates the fraction of TWD1-mCherry fluorescence overlapping with fABD2-GFP fluorescence, respectively. MCC values were calculated as M1 = 0.74 ± 0.035, M2 = 0.71 ± 0.035 (n = 6, high) and M1 = 0.42 ± 0.036, M2 = 0.30 ± 0.031 (n = 6; low). Significant differences (Ordinary one-way ANOVA) ± SE (n = 7) in MCC values are indicated: *, *p* < 0.05; **, *p* < 0.01; ***, *p* < 0.001; ****, *p* < 0.0001. **C:** Interaction of purified TWD1 with rabbit actin (Rb actin) and Arabidopsis Actin2 and Actin7 protein. TWD1 protein was immobilized on magnetic COOH beads and individually incubated with rabbit actin (Rb actin), and purified Arabidopis Actin2 (At actin2) and actin7 (At Actin7; (Kijima et al., 2018)). Bound actin was analyzed by Western blots using monoclonal anti-actin antibodies; empty magnetic beads were used as negative control. Supplementary information can be found in Suppl. Fig. S2.

This is a new and surprising finding, especially in light of the fact that our previous analyses had revealed that interaction between TWD1 and actin is most likely indirect and that the remodeling effect of TWD1 on actin bundling and dynamics thus requires the action of an additional TWD1 interacting protein (Zhu et al., 2016). However, underlying *in vitro* experiments had been performed with rabbit actin only leaving the possibility open that TWD1 might specifically interact with Arabidopsis actin isoforms. Therefore, we repeated actin-pull down experiments with purified TWD1 and Arabidopsis Actin7 protein (Kijima et al., 2018), while using purified Actin2 as negative control. Like with rabbit actin, TWD1 was unable to pull-down Actin7 *in vitro* (Figures 3C). This indicates that TWD1 does most likely not interact directly with actin proteins suggesting the existence of intermediate linker proteins between TWD1 and actin (Zhu and Geisler, 2015; Zhu et al., 2016).

### TWD1 is involved in myosin-dependent ER motility and cargo trafficking

In order to identify such a linker protein that is able to bridge such an indirect TWD1-actin interaction, we re-screened the recently published TWD1 interactome (Zhu et al., 2016) for proteins that have both the potential to execute previously described actin bundling (Zhu et al., 2016) and the here described ER remodeling. Beside verified TWD1 interactors, this short list showed a remarkable enrichment of PM proteins putatively involved in protein trafficking, including several Rab GTPases, DYNAMIN-LIKE3 and a clathrin heavy chain (Zhu et al., 2016). However, myosin XIK (At5g20490; hereafter referred to as XIK), before described to be involved in root hair growth, trichome development, and organelle trafficking (Tominaga et al., 2013), revealed similarly high scores than Actin7 (score, 201.00; emPAI value 0.15; (Zhu et al., 2016)) caught our attention and was thus selected for further analyses.

To further elaborate on the putative TWD1-myosin XIK interaction during ER remodeling, we co-expressed XIK-YFP with the ER marker, ER-CFP, and TWD1-mCherry. A nearly perfect colocalization for both indicated that myosin XIK and TWD1 co-localize on the ER membrane (Figures 4A and 4B). In order to further establish TWD1-myosin XIK interaction, we co-expressed XIK-YFP with TWD1 fused to *Renilla* luciferase (TWD1-Rluc) in tobacco, allowing to quantify interaction by bioluminescence resonance energy transfer (BRET) as described previously (Wang et al., 2013; Hao et al., 2020). Analyses resulted in significant BRET ratios for the XIK-YFP/TWD1-Rluc pair that were twice of the positive control, TWD1-Rluc/ABCB1-YFP (Figures 4C). In summary, co-immunoprecipitation (Zhu et al., 2016), co-immunolocalization and BRET analyses strongly indicate that TWD1 and myosin isoform XIK are physically interacting with each other on the ER.

**Fig. 4:**
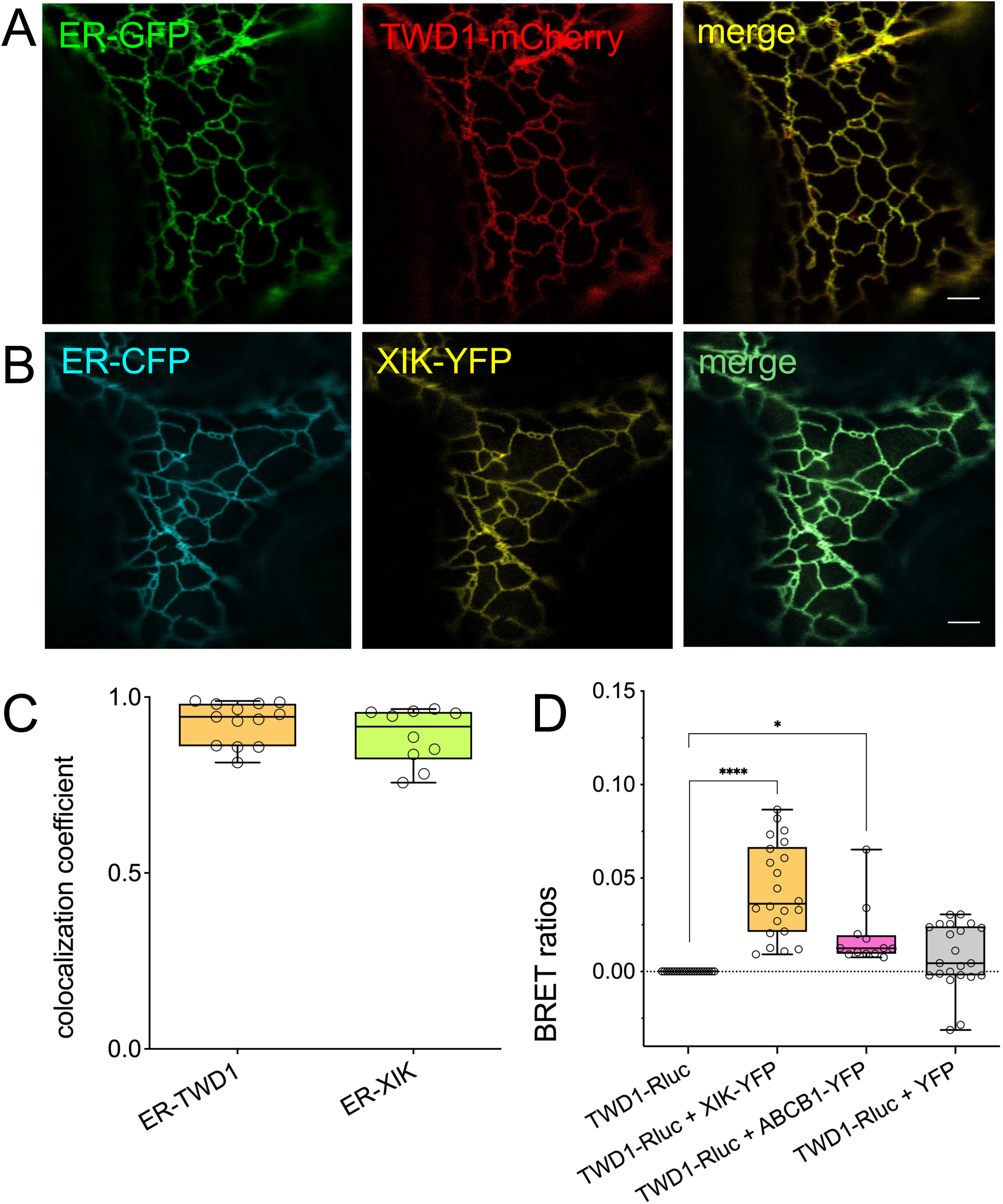
Myosin XI-K functions as a physical linker between actin and TWD1. **A:** ER-GFP (*35S:GFP-HDEL*) and TWD1-mCherry (*35S:TWD1-mCherry*) colocalize when co-expressed in tobacco epidermal leaf cells; bar, 10 μm. **B:** ER-CFP (ER-ck/ CD3-953) and Myosin XIK-YFP (*35S:Myosin XIK-YFP*) colocalize when co-expressed in tobacco epidermal leaf cells; bar, 10 μm. **C:** Quantification of colocalizations in A-B, **D:** Quantification of physical interaction between TWD1-Rluc (*35S:TWD1-Rluc*) and Myosin XIK-YFP assayed by bioluminescence resonance energy transfer (BRET). Tobacco leaves were transfected with *35S:TWD1-Rluc* and *35S:Myosin XIK-YFP* and microsomes were used for BRET measurements; data are means ± SEM (n = 6 independent leaf infiltrations). TWD1-Rluc alone and TWD1-Rluc/YFP were used as negative controls and TWD-Rluc/ABCB1-YFP was used as a positive, respectively (Hao et al., 2020). Significant differences (Ordinary one-way ANOVA) of mean interactions ± SE (n =3 independent tobacco infiltrations) to TWD1-Rluc are indicated: *, *p* < 0.05; **, *p* < 0.01; ***, *p* < 0.001; ****, *p* < 0.0001).

In order to be able to analyze the impact of TWD1 on ER dynamics, we quantified average and maximum velocity flows of ER-localized GFP in WT and *twd1* root epidermal cells as described previously (Ueda et al., 2010; Stefano et al., 2014). The ER strands streamed mainly along the longitudinal axis of the cells (Figure 5A-B, Suppl. Movie S2). Interestingly, ER streaming in *twd1* roots revealed significantly reduced average (0.03 ± 0.006 μm/sec) and maximal velocities (0.93 ± 0.21 μm/sec, respectively, compared to the wild-type (0.07 ± 0.29μm/sec and 1.19 μm/sec ± 0.08 μm/sec, Figure 5C-5D). This strongly suggests that TWD1 is a positive regulator of myosin XIK-dependent ER streaming.

**Fig. 5:**
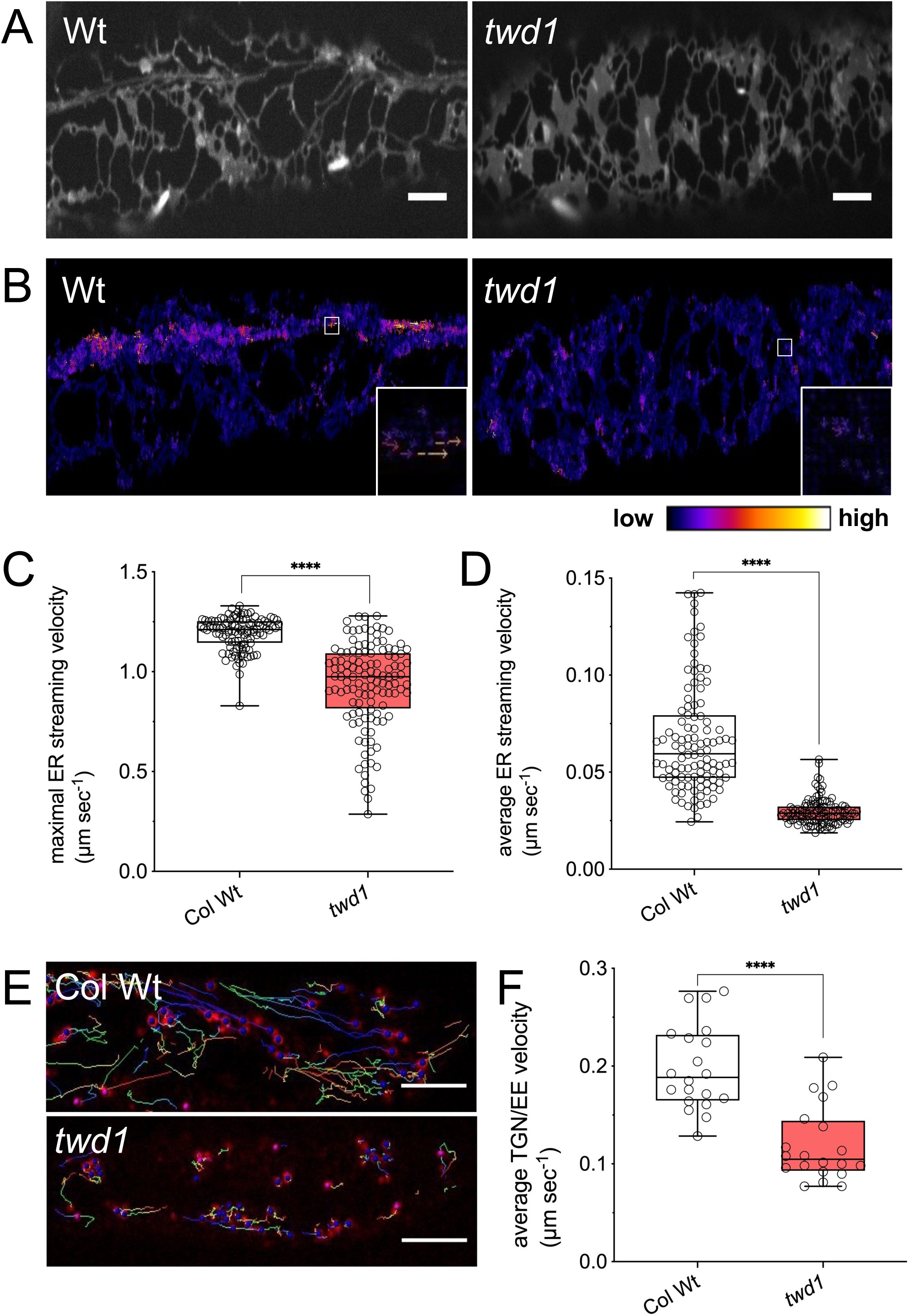
*twd1-3* epidermal root cells have a defect in ER streaming and cargo trafficking. **A:** Confocal spinning-disk analyses of WT (Col-0) and *twd1* (*twd1-3*) 5 dag roots expressing the ER marker, ER-GFP (*35S:GFP-HDEL*). 100 images of GFP-labelled ER at ∼50 ms intervals were captured; bars, 10 μm. Supplementary information can be found in Suppl. Movie S2. **B:** Velocity distribution map generated by processing the time-lapse images of **A**. using LpxFlow (Ueda et al., 2010; Stefano et al., 2014). Arrow lengths and colors indicate streaming velocities; insets are magnified views of the boxed areas in the map. **C-D:** Maximal (**C**) and average velocities (**D**) of ER streaming in Wt (Col-0) and *twd1* (*twd1-3*). Significant differences (unpaired *t* test with Welch’s correction, *p*<0.0001) to Wt are indicated by asterisk; n = 100 cells. **E-F:** Imaging (**E**) and quantification (**F**) of the motility of the TGN/EE marker, VHA-a1-RFP (Dettmer et al., 2006) in Wt and *twd1* (*twd1-3*) using live-cell imaging. Reduced motilities of TGN/EE compartments in *twd1* compared to the wild type (Wt) imply defects in cargo trafficking. Significant differences (unpaired *t* test with Welch’s correction, *p*<0.0001) to Wt are indicated by asterisks; n = 100 cells. Supplementary information can be found in Suppl. Movie S3.

Furthermore, we examined the behavior of the endosomal marker, VHA-a1-RFP, which localizes on the *trans*-golgi network and early endosomes (TGN/EE; (von der Fecht-Bartenbach et al., 2007)) Quantitative analyses of live-cell imaging indicated a significantly reduced motion of TGN/EE compartments in *twd1* compared to the wild type (Figure 5E-F; Suppl. Movie S3), implying that cargo trafficking through these vesicles is inhibited in the absence of TWD1.

### Arabidopsis myosin XI mutants are defective in auxin transport

The findings above suggested that TWD1 positively regulates myosin-dependent ER motility and cargo trafficking, which should have also a direct impact on auxin transporter expression and auxin transport (Abu-Abied et al., 2018). In fact, by using higher-order myosin XI mutants, myosin XI was shown to be involved in root organogenesis and cell division most likely due to defects in polar auxin transport (Abu-Abied et al., 2018). As a proof-of-concept, PIN1 polarity and DR5-GFP reporter expression was reduced in a triple myosin KO line, lacking *XI-K, XI-1* and *XI-2* (*3KO*; (Prokhnevsky et al., 2008; Peremyslov et al., 2010; Abu-Abied et al., 2018), although auxin transport was not quantified.

Using the well-established Arabidopsis protoplast export system (Henrichs et al., 2012; Wang et al., 2013; Hao et al., 2020), we found a gradual reduction of IAA export for single myosin XI mutants, although significant differences to WT were only found with the triple loss-of-function (*3KO*) line (Figure 6A; Suppl. Fig. S3C). This behavior is perfectly in agreement with growth phenotypes (here: petiole lengths; Suppl. Fig. S3A-B), verifying on a transport level the reported functional redundancy between these highly expressed myosins (Prokhnevsky et al., 2008). Reductions in auxin export were restored to WT level in the *3KOR* line expressing *XIK:YFP* in *3KO* (Peremyslov et al., 2012). Specificity of IAA export reductions were further underlined by analyzing the export of benzoic acid (BA), an unspecific diffusion control, that was assayed together with IAA in double labeling experiments (Figure 6B; Suppl. Fig. S3D).

**Fig. 6:**
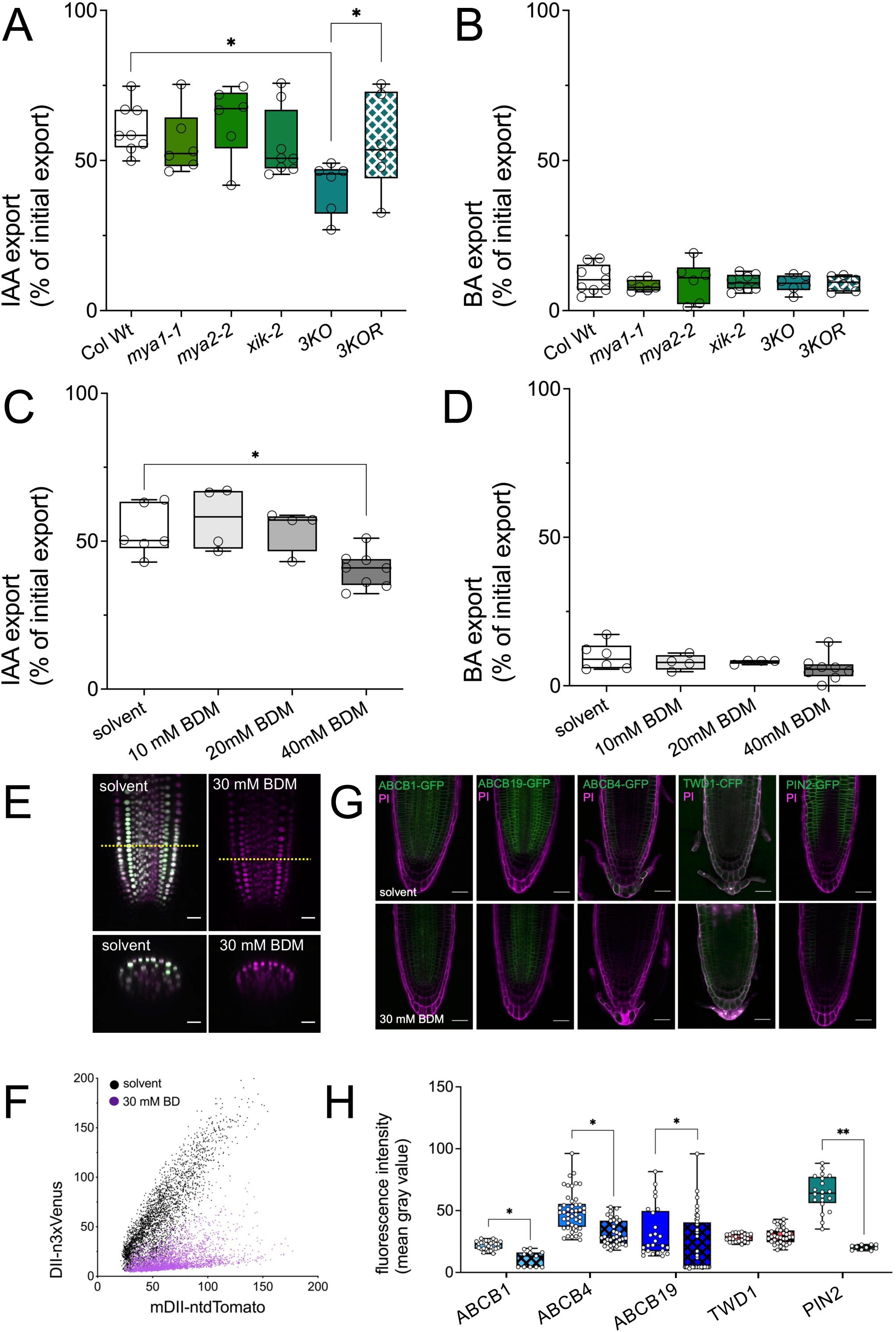
Altering myosin XI presence leads to defects in auxin transport caused by reduced expression of auxin exporters. **A-B**: Reduced IAA (**A**) and benzoic acid (BA; **C**) efflux from Arabidopsis leaf protoplasts prepared from indicated *Myosin XI* loss-of-function alleles. Significant differences (Ordinary one-way ANOVA) ± SE are indicated: *, *p* < 0.05; **, *p* < 0.01; ***, *p* < 0.001; ****, *p* < 0.0001; n≥6 transport experiments generated from independent protoplast preparations. Supplementary information can be found in Suppl. Fig. S3. **C-D:** Reduced IAA (**C**) and benzoic acid (BA; **D**) efflux after BDM treatment of Col Wt (Col-0) plants. Significant differences (Ordinary one-way ANOVA) ± SE are indicated: *, *p* < 0.05; **, *p* < 0.01; ***, *p* < 0.001; ****, *p* < 0.0001; n≥6 transport experiments generated from independent protoplast preparations. Supplementary information can be found in Suppl. Fig. S3. **E-F:** Analyses of the semi-quantitative auxin-input-reporter, R2D2, in wild-type plants treated with 30 mM BDM. mDII-ntdTomato (magenta) and DII-n3xVenus (green) fluorescence signal overlays (**E**) and signal intensities (**F**) in the root apical meristem; bars, 10 μm. Supplementary information can be found in Suppl. Fig. S3. **G-H:** Quantification of ABCB1-GFP, ABCB4-GFP, ABCB19-GFP, TWD1-CFP and PIN2-GFP expression in the Arabidopsis root after treatment with 30mM BDM or solvent. Exemplary confocal images (**G**) and their quantification (**H**); bars, 10 μm. Significant differences (Ordinary one-way ANOVA) of mean expression ± SE (n=6) to solvent control are indicated: *, *p*<0.05; **, *p*<0.01; ***, *p*<0.001; ****, *p*<0.0001).

To further establish myosin XI proteins as relevant actors of auxin export, we treated WT protoplasts with increasing concentrations of 2,3-butanedione monoxime (BDM), a low-affinity inhibitor of skeletal muscle myosins. BDM recently received some criticism in respect to its specificity but that can be safely considered as a general myosin inhibitor (Ostap, 2002). However, increasing concentrations of BDM had the same inhibitory effect on auxin (but not BA) export as the triple loss-of-function mutation (Figure 6C-D; Suppl. Fig. S3E-F). The inhibitory effect of BDM on polar auxin transport was further verified by quantifying Wt roots expression of the semi-quantitative auxin reporter, R2D2 (Liao et al., 2015). 30mM BDM, a concentration below maximal transport inhibition, was sufficient to entirely reduce auxin responses (Figure 6E-F), while lathrunculin B and oryzalin at micro-molar concentration known to disrupt the actin and microtubule cytoskeleton had only mild effects (Suppl. Fig. S3G-H).

To finally provide on one hand a mechanistic explanation for BDM action on auxin transport and to investigate the consequences of myosin inhibition on the other, we analyzed the effect of BDM on ABCB1,4,19-GFP, TWD1-CFP and PIN2-GFP expression in the Arabidopsis root. We found that BDM similarly interfered with the plasma membrane secretion of ABCB-and PIN-type auxin exporters but had no significant effect of TWD1 trafficking and expression (Figure 6G-H). In summary, this data set provides strong indication that myosin XI proteins are required for proper auxin exporter trafficking and thus auxin-mediated plant development.

## DISCUSSION

### TWD1 is essential for actin cytoskeleton organization independent of ABCB-mediated auxin delivery

Defects in auxin transport caused by loss-of TWD1 have so far been accounted to a dual function of TWD1: TWD1 is thought to act on one hand as ABCB chaperone that positively regulates ABCB biogenesis (Wang et al., 2013) and transport activity (Hao et al., 2020) and, on the other, as a modulator of actin cytoskeleton bundling and dynamics by indirect interaction with ACTIN7 (Zhu et al., 2016). However, pinpointing the exact mode of TWD1 action had been hampered by the fact that both pathways are interconnected by the auxin-actin circuit in that altered local auxin concentrations directly or indirectly effect actin conformations, which again has an effect on the positioning of auxin transporters (Zhu and Geisler, 2015; Zhu et al., 2016).

Here, in order to dissect the individual modes of TWD1 we used the unicellular pollen system that unlike so far analyzed hypocotyls and roots is independent of ABCB-mediated auxin provided by neighboring cells. Moreover, *ABCB1,4,19* expression known to be controlled by TWD1 is neglectable in mature pollen (Suppl. Fig. S1A), excluding also intracellular auxin gradients eventually provided by ABCBs. Our analyses revealed that TWD1 is increasingly expressed in germinating pollen and pollen tubes and positively regulates pollen tube physiology characterized by pollen germination and growth (Figure 1; Suppl. Fig. S1; Suppl. Movie S1). Importantly, these defects are in contrast to most described developmental phenotypes found for *twd1* not phenocopied in *abcb1 abcb19* indicating an ABCB-independent action. Interestingly, TWD1 changes its expression profile with pollen germination to a ‘collar-like’ structure at the site of germination (Figure 1; Suppl. Fig. S1; Suppl. Movie S1) resembling the actin cytoskeleton at the site of pollen germination (Pierson and Cresti, 1992; Gibbon et al., 1999; Wu et al., 2010; Liu et al., 2018). After germination, the TWD1-CFP signal was found to be concentrated in the subapical region of the pollen tube (Suppl. Movie S1) that contains a highly dynamic “actin fringe” structure as well as a peak region for exocytosis.

This functional correlation between TWD1 expression and actin cytoskeleton organization during pollen germination was hardened by the finding that *TWD1*-loss-of-function alters actin organization and tip-directed vesicle transport in pollen tubes (Figure 2). Our data thus provide evidence that TWD1 plays a key role in pollen germination and pollen tube tip growth by controlling actin organization, however, importantly in an action independent of ABCB1,19-mediated auxin delivery indicating a direct impact of TWD1 on actin remodeling.

### Myosin XI-K functions as a physical linker between actin and TWD1 and is putatively involved in actin re-organization

Our previous work established a role of TWD1 in actin remodeling most likely by physical interaction between TWD1 and ACTIN7 that, however, based on pull-down studies with mammalian F-actin were considered as being of indirect nature (Zhu and Geisler, 2015; Zhu et al., 2016). Using ultra-pure Arabidopsis TWD1 and ACTIN7 protein, this indirect interaction could be re-confirmed (Figure 3; Suppl. Figure S2D). Interestingly, by BRET analyses we verified with myosin XI-K a recently identified interacting protein as valid TWD1 interactor (Figure 4) functioning most likely as a physical linker between actin and TWD1. Interestingly, myosin XI-K does not only perfectly co-localize with TWD1, a predominant ER constituent, but also with the established ER marker, ER-CFP (ER-ck/ CD3-953 (Nelson et al., 2007); Figure 4).

Several studies have indicated that myosin XI acts as a direct regulator of F-actin organization (Duan and Tominaga 2018): treatment with the myosin ATPase inhibitor, BDM, results in disorganization of F-actin in root hairs, and moreover, F-actin dynamics were drastically inhibited in the myosin 3KO mutant (Tominaga et al. 2000). Our results thus suggest that myosin XI-K not only bridges TWD1-actin interaction but also most likely conduct actin bundling on the actin cytoskeleton previously diagnosed (Zhu et al., 2016; Tan et al., 2020).

Recently, myosin XIs were shown to be involved in Arabidopsis development caused most likely by defects in the polar distribution of auxin (Ojangu et al. 2018, Abu-Abied et al. 2018). Here, we verify that myosin XI mutants are indeed defective in auxin transport and that exact three myosin XI isoforms that showed overlapping function in cytoplasmic streaming and organelle trafficking (Peremyslov et al., 2008, 2010; Prokhnevsky et al., 2008; Ueda et al., 2010) functional redundantly in respect to auxin efflux (Figure 6; Suppl. Fig. S3). Interestingly, the roughly 30% reduction in auxin efflux for the *3KO* mutant was very similar to what was previously found for the *TWD1* loss-of-function allele (Bouchard et al., 2006) and comparable with the pharmacological inhibition of all myosins by BDM (Figure 6). This suggests that the TWD1-myosinXI-actin module is primarily responsible for (ABCB-mediated) auxin efflux controlled by TWD1.

An interesting, though puzzling finding was that based on BDM treatments (Figure 6) and genetics (*3KO*; Abu-Abied et al. 2018) PIN and ABCB-type auxin exporters were delocalized in myosin XI mutants, as was found for *actin7* (Zhu et al., 2016; Abu-Abied et al., 2018), while apparently in *twd1* only a few auxin-transporting ABCBs but not other auxin exporters (or PM-intrinsic proteins) are affected in their biogenesis (Wu et al., 2010; Wang et al., 2013). Although one has to be generally be careful with the interpretation of inhibitor studies, these data might suggest that actually TWD1 provides specificity toward ABCB secretion via the TWD1-myosinXI-actin module.

In summary, these data sets provide strong indication that myosin XI proteins are required for proper auxin exporter trafficking and thus auxin export by linking and conducting the action of TWD1 on actin cytoskeleton remodeling.

### TWD1 is an ER–actin adapter proteins involved in myosin-dependent ER motility and cargo trafficking

The dynamic three-way interactions between ER, F-actin and myosin XIs does not only determine the architecture and movement of thick ER strands and is the primary cause of cytoplasmic streaming, but also connects the ER with cortical F-actin. However, essential linker or adapter proteins that facilitate and/or regulate anchoring of the ER membrane with the actin cytoskeleton are hardly known. Our previous and current data qualify the FKBP42, TWD1, as such a functional ER–actin adapter: First, TWD1 is a prominent ER resident that is attached most likely by in-plane membrane anchoring to the outer surface of the ER membrane (Scheidt et al., 2006). Second, TWD1 physically binds both myosin XIK (Figure 6) and F-actin (Zhu et al., 2016), the latter most likely indirectly via myosin XIK (Zhu et al., 2016; Fig. 3; Suppl. Fig. S2). Third, TWD1 controls both actin bundling and dynamics (Fig. 2; Zhu and Geisler, 2015; Zhu et al., 2016; Tan et al., 2020), as well as ER motility and vesicle trafficking (Figure 5). Strikingly, and fourth, the ER-localized TWD1 reshapes the ER network to overlay actin cables (Fig. 3; Suppl. Fig. S2) similar to myosin-XI (Ueda et al., 2010). This resembles strongly the action of the SNARE protein, SYP73, where likewise *SYP73* over-expression caused a rearrangement of the ER over actin and *SYP73* loss of-function reduced ER streaming and plant growth (Cao et al., 2016). The current picture that emerges is that ER–actin adapter proteins, like TWD1 (and SYP73), modulate ER structure and movement by working in conjunction with myosin XI proteins. The adapter proteins would bring ER and F-actin closer together allowing the myosin driven movement to take place resulting in rearrangement of the F-actin. However, a striking difference is that SYP73 contains actin binding domains and is an actin-binding protein itself (Cao et al., 2016) suggesting an overall different mode of assembly. TWD1 as an ER-actin adaptor is at first view surprising, as so far only actins, myosin XIs and ER-related proteins were reported as being involved in cytoplasmic streaming. However, support for a direct regulatory involvement of a component of the auxin efflux machinery in this process was provided by the finding that cytoplasmic steaming was drastically impaired by depletion of *ABCB19* (Okamoto et al., 2016).

In conclusion, our data identify TWD1 as a novel myosin XIK interacting protein that is required for actin-dependent plant growth independent of its action as an ABCB chaperone. TWD1 seems to bridge the ER with actin and has an important role in ER shaping and vesicle cargo streaming in conjunction with the actin-myosin XI module. As such our findings provide a molecular explanation for the defects in early ABCB biogenesis that cause a major part of the pleiotropic *twisted dwarf* syndrome

## MATERIALS AND METHODS

### Accession numbers

Sequence data from this article can be found in the Arabidopsis Genome Initiative or GenBank/EMBL databases under the following accession numbers: *TWD1*, At3g21640; *ABCB1* (formerly *PGP1*), At2g36910; *ABCB4* (formerly *PGP4*), At2g47000; *ABCB19* (formerly *MDR1* or *PGP19*), At3g28860; *MYA1* (formerly *MYOSIN XI-1*), At1g17580; *MYA2* (formerly *MYOSIN XI-2*), At5g43900; *XI-K* (*formerly XI-17*), At5g20490

### Growth conditions and sample collection

Arabidopsis plants were grown for 15 days at 22°C under an 8h light/16h dark cycle and then transferred to a 16h/8h dark cycle until flowering, pollen was collected one week after first flowering. To quantify germination rates, pollen grains from mutants and wild type were cultures on the same medium for 3h/ 6h and observed using a microscope under brightfield (DM4000B, Leica; Liu et al., 2018).

### Time-Lapse Imaging and Image Analysis

To visualize TWD1-CFP, the vesicle marker, Rab4d-YFP (*35S: Rab4d-YFP*; (Lee et al., 2008), and actin filament dynamics during pollen germination, pollen grains were cultures for 2h before microscopy. Time-lapse imaging was performed using a spinning-disk confocal microscope (Visitron VisiScope CSU-W1) and 488-and 561-nm lasers. Z-stack time-series were collected at maximal speed with the z-step set at 0.5 mm.

To quantify ER streaming velocities, five-day-old roots of transgenic plants expressing ER-GFP (*35S:GFP-HDEL;* Mitsuhashi et al., 2000) were observed by spinning-disk confocal microscopy (Visitron VisiScope CSU-W1). 100 images of GFP-labelled ER at ∼50 ms intervals for each cell were captured. Velocity and velocity distribution maps were generated by processing the time-lapse images using the LpxFlow ImageJ plug-in (https://lpixel.net/en/products/lpixel-imagej-plugins) (Ueda et al., 2010)). To quantify co-localization of TWD1-mCherry with the ER marker, ER-GFP (*35S:GFP-HDEL*), and actin markers, fABD2-GFP or Lifeact-GFP (Zhu et al., 2016), the co-localization plugin for ImageJ to calculate Manders split coefficients and Pearson’s correlation coefficient for red and green signals was used. For confocal laser scanning microscopy work, the Leica TCS-SP5 was used. Confocal settings were set to record the emission of GFP (excitation 488 nm; emission 505 to 530 nm), CFP (excitation 458 nm; emission 465 to 500 nm), chloroplast autofluorescence (excitation 488 nm; emission 650 to 710 nm). Images were acquired with the ImageJ software (http://imagej.nih.gov/ij/) using identical settings for all samples.

### RNA Extraction and qRT–PCR

Hundreds of fresh flowers were harvested to obtain pollen grains. Total RNA from pollen grains was isolated with TRIzol reagent (Invitrogen) according to the manufacturer’s instructions. The cDNA was reverse-transcribed from total RNA using Omniscript® reverse transcriptase (QIAGEN). The expression levels of *TWD1* were determined via qRT–PCR using the SYBR Green mix, and UBQ10 was amplified as an internal control.

### *In vitro* actin binding assays

Purified TWD1 protein (Zhu et al., 2016) was immobilized on activated magnetic COOH beads at room temperature. TWD1 (or empty) beads were washed with PBS pH 7.4 and blocked with 0.1M Tris pH7.0. TWD1-beads were incubated with rabbit actin (Rb actin), purified Arabidopsis Actin2 (At Actin2) or actin7 (At Actin7; (Kijima et al., 2018)) for 2h at room temperature, respectively. Bound actin was eluted with 0.1 M Glycin pH 2.8 and analyzed by Western blots using monoclonal anti-actin antibodies; empty magnetic beads were used as negative control.

Prior to use, G-actin and GST-TWD1 expressed and purified from *E. coli* BL21(DE3) were centrifuged at 55,000 rpm, 4°C for 60 min. and the supernatant was used. G-actin was polymerized at 25°C for 30 min in 10x KMEI buffer (500 mM KCl, 10 mM MgCl_2_, 10 mM EGTA, and 100 mM imidazole-HCl, pH 7.0). Polymerized F-actin (3 μM) was incubated with various concentrations (0, 0.25, 0.5, 1, 2, 3, 6 μM) of GST-TWD1 on ice for 30 min. After centrifugation at 40,000 rpm at 4°C for 30 min, collected supernatants and pellets were analyzed by SDS-PAGE.

### Myosin-TWD1 interaction analyses

For BRET analysis, *N. benthamiana* leaves were Agrobacterium co-infiltrated with indicated BRET construct combinations (or corresponding empty vector controls) and microsomal fractions were prepared 4days after inoculation (dai). BRET signals were recorded from microsomes (each ∼10 μg) in the presence of 5 µM coelenterazine (Biotium Corp.) using the Cytation 5 image reader (BioTek Instruments) and BRET ratios were calculated as described previously (Wang et al., 2013). The results are the average of 20 readings collected every 30 seconds, presented as average values from a minimum of three independent experiments (biological replica: independent Agrobacterium infiltrations and microsome preparations) each with four technical replicates.

### Auxin transport

Simultaneous ^3^H-3-indolylacetic acid (IAA) and ^14^C-benzoic acid (BA) export from Arabidopsis mesophyll protoplasts was analysed as described (Henrichs et al., 2012). Relative export from protoplasts was calculated from exported radioactivity into the supernatant as follows: (radioactivity in the supernatant at time t = x min.) - (radioactivity in the supernatant at time t = 0)) * (100%)/ (radioactivity in the supernatant at t = 0 min.); presented are mean values from >4 independent protoplast preparations.

### Statistical analysis

All statistical analyses were performed using Prism 9.3.1 (GraphPad Software). Confocal images were processed in Adobe Photoshop 2020 (Adobe Systems) and quantified using ImageJ software (http://imagej.nih.gov/ij/).

## ACKNOWLEDGMENTS

We thank Haruko Ueda (Konan University, Japan) for providing us with the Myosin XI-deficient mutants and the protocol using LpxFlow software analysis the ER streaming, Taro Uyeda (Waseda University, Japan) for purified ACT2 and ACT7 protein, Jiri Friml (Institute of Science and Technology Austria, Austria) for VHA-a1-RFP transgenic lines in wild-type and *twd1-3* and Zhenbiao Yang (University of California at Riverside, USA) for Rab4d-YFP plasmid and transgenic lines of Rab4d-YFP. This work was supported by Swiss National Funds (project 31003A_165877 and 310030_197563) to M.G.

## AUTHOR CONTRIBUTIONS

MG and JL designed the project and JL performed most experiments. PH, JZ and MdD helped with BRET, pollen and molecular work, respectively. JL, JZ, MdD and PH were supervised by MG, while JL was also supervised by HR. Data were analyzed by JL, JZ, PH and MG and MG wrote the manuscript; all authors commented on the manuscript.

## Supplemental data

**The following materials are available in the online version of this article:**

**Suppl. Fig. S1: TWD1 is expressed in germinating pollen and pollen tubes**

**A:** RNA-seq analyses of *TWD1* and selected auxin transporter genes in Arabidopsis pollen and flower taken from a public-available database (http://travadb.org).

**B-C:** The TWD1-CFP labels a “collar-like” structure at the pollen germination site

(**B**). (**C**) Three-dimensional heatmap rendering of the fluorescence intensity of TWD1-CFP (rectangle in **B**).

**Suppl. Fig. S2: TWD1 is localized at the ER and does not directly interact with actin**

**A:** TWD1-mCherry expression overlaps with the ER marker, ER-GFP.

**B:** Co-expression of TWD1-mCherry with microtubule marker, TUA6-GFP, indicates that the modified ER network labeled by TWD1-mCherry does not overlap with the microtubule cytoskeleton.

**C:** High expression of PIN6-GFP with the ER lumen marker, ERYK, did not lead to ER rearrangement to actin-like organization supporting the specificity of the ER shaping effect of TWD1.

**Suppl. Fig. S3: Arabidopsis myosin XI mutants are defective in auxin transport A-B**: Growth phenotypes (**A**) and quantification of petiole lengths (**B**) of indicated *Myosin XI* loss-of-function plants. Significant differences (unpaired *t* test with Welch’s correction, *p*<0.05) are indicated by asterisks (mean ± SEM; n ≥ 6).

**C-D:** Time kinetics of IAA (**C**) and benzoic acid (BA) efflux (**D**) from leaf protoplasts prepared from indicated *Myosin XI* loss-of-function plants (mean ± SEM; n ≥ 6 transport experiments generated from independent protoplast preparations).

**E-F:** Time kinetics of IAA (**E**) and benzoic acid (BA) efflux (**F**) from leaf protoplasts prepared from Wt after BDM treatment (mean ± SEM; n ≥ 6 transport experiments generated from independent protoplast preparations).

**G-H:** Imaging (**G**) and quantification (**H**) of the semiquantitative auxin-input-reporter, R2D2, after 12h treatment with Lathrunculin B (LatB; 1μM) and Oryzalin (25μM) treatment; scale bar, 10 μm

**Suppl. Movie S1: The subcellular localization of TWD1 changes during pollen tube germination and elongation**.

Pollen grains expressing TWD1:TWD1-CFP in the *twd1-1* background were germinated *in vitro* and observed using spinning disc confocal microscopy. Before germination, TWD1-GFP is detected in the cytoplasm but later subsequently localized to a “collar-like” structure at the germination site. Images were captured every 1 min for 15 min; bar = 10 mm.

**Suppl. Movie S2: Expression of the ER marker, ER-GFP (*35S:ER-GFP*), in the Arabidopsis Wt and *twd1* (*twd1-3*) root**.

Confocal images of root epidermal cells of 5DAG wild-type (Col-0) and *twd1-3* expressing the ER marker ERYK; bars, 10 μm.

**Suppl. Movie S3: Motility of the TGN/EE marker, VHA-a1-RFP, in Wt and *twd1***. Live-cell imaging of root epidermal cells of 5 DAG wild-type (Col-0) and *twd1-3* expressing the TGN/EE marker, VHA-a1-RFP; bars, 10 μm.

## REFERENCES

Abu-Abied M, Belausov E, Hagay S, Peremyslov V, Dolja V, Sadot E (2018) Myosin XI-K is involved in root organogenesis, polar auxin transport, and cell division. J Exp Bot 69: 2869–2881

Bouchard R, Bailly A, Blakeslee JJ, Oehring SC, Vincenzetti V, Lee OR, Paponov I, Palme K, Mancuso S, Murphy AS, Schulz B, Geisler M (2006) Immunophilin-like TWISTED DWARF1 modulates auxin efflux activities of Arabidopsis P-glycoproteins. J Biol Chem 281: 30603–30612

Cao P, Renna L, Stefano G, Brandizzi F (2016) SYP73 Anchors the ER to the Actin Cytoskeleton for Maintenance of ER Integrity and Streaming in Arabidopsis. Curr Biol 26: 3245–3254

Corti B (1774) Osservazioni microscopiche sulla tremella e sulla circolazione del fluido in una pianta acquajuola. .

Dettmer J, Hong-Hermesdorf A, Stierhof YD, Schumacher K (2006) Vacuolar H+-ATPase activity is required for endocytic and secretory trafficking in Arabidopsis. Plant Cell 18: 715–730

Dhonukshe P, Grigoriev I, Fischer R, Tominaga M, Robinson DG, Hasek J, Paciorek T, Petrasek J, Seifertova D, Tejos R, Meisel LA, Zazimalova E, Gadella TW, Jr., Stierhof YD, Ueda T, Oiwa K, Akhmanova A, Brock R, Spang A, Friml J (2008) Auxin transport inhibitors impair vesicle motility and actin cytoskeleton dynamics in diverse eukaryotes. Proc Natl Acad Sci U S A 105: 4489–4494

Duan Z, Tominaga M (2018) Actin-myosin XI: an intracellular control network in plants. Biochem Biophys Res Commun 506: 403–408

Feraru E, Vosolsobe S, Feraru MI, Petrasek J, Kleine-Vehn J (2012) Evolution and Structural Diversification of PILS Putative Auxin Carriers in Plants. Front Plant Sci 3: 227

Geisler M, Hegedus T (2020) A twist in the ABC: regulation of ABC transporter trafficking and transport by FK506-binding proteins. FEBS Lett 594: 3986–4000

Geisler M, Kolukisaoglu HU, Bouchard R, Billion K, Berger J, Saal B, Frangne N, Koncz-Kalman Z, Koncz C, Dudler R, Blakeslee JJ, Murphy AS, Martinoia E, Schulz B (2003) TWISTED DWARF1, a unique plasma membrane-anchored immunophilin-like protein, interacts with Arabidopsis multidrug resistance-like transporters AtPGP1 and AtPGP19. Mol Biol Cell 14: 4238–4249

Gibbon BC, Kovar DR, Staiger CJ (1999) Latrunculin B has different effects on pollen germination and tube growth. Plant Cell 11: 2349–2363

Hao P, Xia J, Liu J, Di Donato M, Pakula K, Bailly A, Jasinski M, Geisler M (2020) Auxin-transporting ABC transporters are defined by a conserved D/E-P motif regulated by a prolylisomerase. J Biol Chem 295: 13094–13105

Henrichs S, Wang B, Fukao Y, Zhu J, Charrier L, Bailly A, Oehring SC, Linnert M, Weiwad M, Endler A, Nanni P, Pollmann S, Mancuso S, Schulz A, Geisler M (2012) Regulation of ABCB1/PGP1-catalysed auxin transport by linker phosphorylation. EMBO J 31: 2965–2980

Kijima ST, Staiger CJ, Katoh K, Nagasaki A, Ito K, Uyeda TQP (2018) Arabidopsis vegetative actin isoforms, AtACT2 and AtACT7, generate distinct filament arrays in living plant cells. Sci Rep 8: 4381

Lee YJ, Szumlanski A, Nielsen E, Yang Z (2008) Rho-GTPase-dependent filamentous actin dynamics coordinate vesicle targeting and exocytosis during tip growth. J Cell Biol 181: 1155–1168

Liao CY, Smet W, Brunoud G, Yoshida S, Vernoux T, Weijers D (2015) Reporters for sensitive and quantitative measurement of auxin response. Nat Methods 12: 207–210, 202 p following 210

Liu C, Zhang Y, Ren H (2018) Actin Polymerization Mediated by AtFH5 Directs the Polarity Establishment and Vesicle Trafficking for Pollen Germination in Arabidopsis. Mol Plant 11: 1389–1399

Liu J, Ghelli R, Cardarelli M, Geisler M (2022) TWISTED DWARF1 regulates Arabidopsis stamen elongation by differential activation of ABCB1,19-mediated auxin transport. J Exp Bot

Lovy-Wheeler A, Wilsen KL, Baskin TI, Hepler PK (2005) Enhanced fixation reveals the apical cortical fringe of actin filaments as a consistent feature of the pollen tube. Planta 221: 95–104

Mao H, Nakamura M, Viotti C, Grebe M (2016) A Framework for Lateral Membrane Trafficking and Polar Tethering of the PEN3 ATP-Binding Cassette Transporter. Plant Physiol 172: 2245–2260

Mitsuhashi N, Shimada T, Mano S, Nishimura M, Hara-Nishimura I (2000) Characterization of organelles in the vacuolar-sorting pathway by visualization with GFP in tobacco BY-2 cells. Plant Cell Physiol 41: 993–1001

Montes-Rodriguez A, Kost B (2017) Direct Comparison of the Performance of Commonly Employed In Vivo F-actin Markers (Lifeact-YFP, YFP-mTn and YFP-FABD2) in Tobacco Pollen Tubes. Front Plant Sci 8: 1349

Nelson BK, Cai X, Nebenfuhr A (2007) A multicolored set of in vivo organelle markers for co-localization studies in Arabidopsis and other plants. Plant J 51: 1126–1136

Okamoto K, Ueda H, Shimada T, Tamura K, Koumoto Y, Tasaka M, Morita MT, Hara-Nishimura I (2016) An ABC transporter B family protein, ABCB19, is required for cytoplasmic streaming and gravitropism of the inflorescence stems. Plant Signal Behav 11: e1010947

Ostap EM (2002) 2,3-Butanedione monoxime (BDM) as a myosin inhibitor. J Muscle Res Cell Motil 23: 305–308

Peremyslov VV, Klocko AL, Fowler JE, Dolja VV (2012) Arabidopsis Myosin XI-K Localizes to the Motile Endomembrane Vesicles Associated with F-actin. Front Plant Sci 3: 184

Peremyslov VV, Prokhnevsky AI, Dolja VV (2010) Class XI myosins are required for development, cell expansion, and F-Actin organization in Arabidopsis. Plant Cell 22: 1883–1897

Pierson ES, Cresti MJIRoC (1992) Cytoskeleton and cytoplasmic organization of pollen and pollen tubes. 140: 73–125

Prokhnevsky AI, Peremyslov VV, Dolja VV (2008) Overlapping functions of the four class XI myosins in Arabidopsis growth, root hair elongation, and organelle motility. Proc Natl Acad Sci U S A 105: 19744–19749

Riedl J, Crevenna AH, Kessenbrock K, Yu JH, Neukirchen D, Bista M, Bradke F, Jenne D, Holak TA, Werb Z, Sixt M, Wedlich-Soldner R (2008) Lifeact: a versatile marker to visualize F-actin. Nat Methods 5: 605–607

Scheidt HA, Vogel A, Eckhoff A, Koenig BW, Huster D (2006) Solid-state NMR characterization of the putative membrane anchor of TWD1 from Arabidopsis thaliana. Eur Biophys J

Shimmen T, Yokota E (2004) Cytoplasmic streaming in plants. Curr Opin Cell Biol 16: 68–72

Sparkes I, Runions J, Hawes C, Griffing L (2009) Movement and remodeling of the endoplasmic reticulum in nondividing cells of tobacco leaves. Plant Cell 21: 3937–3949

Sparkes IA, Frigerio L, Tolley N, Hawes C (2009) The plant endoplasmic reticulum: a cell-wide web. Biochem J 423: 145–155

Stefano G, Renna L, Brandizzi F (2014) The endoplasmic reticulum exerts control over organelle streaming during cell expansion. J Cell Sci 127: 947–953

Sweeney BM, Thimann KV (1942) The Effect of Auxins on Protoplasmic Streaming. Iii. J Gen Physiol 25: 841–854

Tan S, Di Donato M, Glanc M, Zhang X, Klima P, Liu J, Bailly A, Ferro N, Petrasek J, Geisler M, Friml J (2020) Non-steroidal Anti-inflammatory Drugs Target TWISTED DWARF1-Regulated Actin Dynamics and Auxin Transport-Mediated Plant Development. Cell Rep 33: 108463

Tominaga M, Kimura A, Yokota E, Haraguchi T, Shimmen T, Yamamoto K, Nakano A, Ito K (2013) Cytoplasmic streaming velocity as a plant size determinant. Dev Cell 27: 345–352

Twell D, Wing R, Yamaguchi J, McCormick S (1989) Isolation and expression of an anther-specific gene from tomato. Mol Gen Genet 217: 240–245

Ueda H, Tamura K, Hara-Nishimura I (2015) Functions of plant-specific myosin XI: from intracellular motility to plant postures. Curr Opin Plant Biol 28: 30–38

Ueda H, Yokota E, Kutsuna N, Shimada T, Tamura K, Shimmen T, Hasezawa S, Dolja VV, Hara-Nishimura I (2010) Myosin-dependent endoplasmic reticulum motility and F-actin organization in plant cells. Proc Natl Acad Sci U S A 107: 6894–6899

von der Fecht-Bartenbach J, Bogner M, Krebs M, Stierhof YD, Schumacher K, Ludewig U (2007) Function of the anion transporter AtCLC-d in the trans-Golgi network. Plant J 50: 466–474

Wang B, Bailly A, Zwiewka M, Henrichs S, Azzarello E, Mancuso S, Maeshima M, Friml J, Schulz A, Geisler M (2013) Arabidopsis TWISTED DWARF1 functionally interacts with auxin exporter ABCB1 on the root plasma membrane. Plant Cell 25: 202–214

Wang P, Hussey PJ (2017) NETWORKED 3B: a novel protein in the actin cytoskeleton-endoplasmic reticulum interaction. J Exp Bot 68: 1441–1450

Wu G, Otegui MS, Spalding EP (2010) The ER-localized TWD1 immunophilin is necessary for localization of multidrug resistance-like proteins required for polar auxin transport in Arabidopsis roots. Plant Cell 22: 3295–3304

Wu Y, Yan J, Zhang R, Qu X, Ren S, Chen N, Huang S (2010) Arabidopsis FIMBRIN5, an actin bundling factor, is required for pollen germination and pollen tube growth. Plant Cell 22: 3745–3763

Ye J, Zheng Y, Yan A, Chen N, Wang Z, Huang S, Yang Z (2009) Arabidopsis formin3 directs the formation of actin cables and polarized growth in pollen tubes. Plant Cell 21: 3868–3884

Zhu J, Bailly A, Zwiewka M, Sovero V, Di Donato M, Ge P, Oehri J, Aryal B, Hao P, Linnert M, Burgardt NI, Lucke C, Weiwad M, Michel M, Weiergraber OH, Pollmann S, Azzarello E, Mancuso S, Ferro N, Fukao Y, Hoffmann C, Wedlich-Soldner R, Friml J, Thomas C, Geisler M (2016) TWISTED DWARF1 Mediates the Action of Auxin Transport Inhibitors on Actin Cytoskeleton Dynamics. Plant Cell 28: 930–948

Zhu J, Geisler M (2015) Keeping it all together: auxin-actin crosstalk in plant development. J Exp Bot 66: 4983–4998

